# Directional information flow as a tool for analyzing protein allostery

**DOI:** 10.64898/2026.06.22.733418

**Authors:** Remy A. Yovanno, Albert Y. Lau

## Abstract

The ability to tune protein function through the binding of modulatory ligands enables the development of therapeutics that steer a biological system away from dysfunctional states underlying disease. Understanding the dynamic mechanisms by which allosteric ligands alter protein function remains an important open question.

Dynamical network models allow us to quantify information flow between protein functional sites. However, existing network models use time-symmetric metrics for computing information from correlated residue motions extracted from molecular dynamics (MD) simulations, failing to fully capture directional information flow between sites. Here, we developed a Python library, *TEntroPy*, and analysis workflow using transfer entropy to generate a directional protein network from equilibrium MD trajectories. Applying this workflow to proteins with known allosteric ligands, we identified residues in both allosteric and orthosteric (primary) binding sites acting as broadcasters and receivers of information. We then computed optimal paths of directional information flow between binding sites. The presence of temporal asymmetry in residue coupling identified from simulations of the unbound (apo) state suggests that directional information flow is encoded in the intrinsic dynamics of the protein. To test this, we perturbed key binding-site residues and demonstrated that our TE-weighted network captures perturbation-induced changes in dynamics along communication routes between binding sites. Our work highlights transfer entropy as an additional tool for understanding the changes in flexibility and rigidity that drive dynamic mechanisms of allosteric communication.

## INTRODUCTION

Underlying protein allostery is the flow of information between important functional sites. Ligand binding perturbs protein dynamics that alter behavior at distant sites. Additionally, changes in binding-site dynamics alter the affinity with which ligands bind. This interplay between ligand binding and protein dynamics is the foundation of allostery, which relies on pathways of communication throughout proteins, explaining how a change at one site can be “felt” at another.

Communication in proteins relies on a network of interacting residues through which information can flow. Methods such as normal mode analysis (NMA)^1^ and Gaussian network models (GNMs)^2,3^ have been applied to proteins to predict dynamic properties from protein structures. While more computationally expensive, molecular dynamics simulations provide additional insight by sampling protein dynamics in a time-resolved manner. Dynamical network models use molecular dynamics trajectories to construct a contact map of interacting residues and quantify the strength of these interactions as the amount of correlation between residue fluctuations^4,5^. From these dynamical networks, we can compute important graph properties such as communities, betweenness, eigenvector centrality, and optimal paths^4,5^.

In dynamical network models, correlations between residues represent the probability of information transfer between them^4^. While the original implementation of these models quantifies inter-residue coupling using the Pearson correlation coefficient, a newer implementation using mutual information has been used to capture non-linear relationships between residue motions^5,6^. In recent years, computing mutual information from molecular dynamics simulations has been used to assess inter-residue coupling^7–10^ and even generate order parameters for enhanced sampling^11^.

Although mutual information has been useful for quantifying information flow in proteins, computing correlations based on mutual information only captures time-symmetric relationships between variables of interest, which do not provide information about the direction of information flow. The time-dependent information metric of transfer entropy (TE)^12^ has been applied to molecular dynamics simulations^13–15^ and GNMs^16–18^ to analyze inter-residue coupling. Transfer entropy has also been used to determine directional relationships in native contact analysis^19^ and glycan conformational dynamics^20^. A related statistical approach, Granger causality^21^, has been applied to molecular timeseries data to quantify temporal relationships from linear couplings between atoms^22–24^. These approaches rely on directional information flow to develop a causal understanding of protein dynamics. Transfer entropy computed from molecular dynamics simulations can be compared to experimental approaches for measuring information flow in proteins^25^, such as UV resonance Raman spectroscopy^2,26–28^, time-resolved infrared spectroscopy^29,30^, and NMR relaxation dispersion experiments^31^.

Results of previous studies applying transfer entropy to molecular dynamics trajectories suggests that these temporal relationships are an intrinsic property of protein dynamics^15^. Additionally, a survey study using GNMs of known allosterically regulated proteins revealed correlation between allosteric and orthosteric ligand-binding sites^32^. Therefore, combining these insights and incorporating a dynamical network may inform whether a network of directional information flow emerging from apo protein dynamics would allow us to predict the effects of a drug binding. Specifically, this network would inform which residues transmit allosteric signals and the paths of information flow connecting important functional sites. However, to address questions of this causal nature requires the development of new tools to incorporate transfer entropy into a dynamical network workflow and extract key properties from a directional protein network.

Here, we introduce a Python library *TEntroPy* and an analysis workflow that uses transfer entropy to quantify directional information flow between pairs of residues and construct a causal protein network from molecular dynamics simulations. From this directional network, we identified residues that transmit allosteric signals and determined how those residues are related to one another in the network by calculating optimal paths between binding sites. These insights served as predictions for how the network should respond to perturbation. We tested these predictions by applying restraints to mimic ligand binding and measuring the response of residues along optimal paths between binding sites. Our analyses of checkpoint kinase 1 (Chk1), TEM-1 *β*- lactamase (TEM), and cyclin-dependent kinase 2 (CDK2) demonstrate the value of transfer entropy as a tool to aid drug discovery efforts.

## METHODS

### Constructing a protein network

The *TEntroPy* Python package requires the input of a molecular dynamics (MD) trajectory and a protein contact map for all pairs of residues where residues in contact are denoted with a 1 and those not in contact are denoted with a 0. This contact map can be generated any number of ways, but here, contact maps were generated using the dynamical network analysis Python package dynetan with a stride of 100 ps (10 windows for 1 *μ*s, 5 for 500 ns), as demonstrated in previous work^5^. In this protocol, residues are considered in contact if their heavy atoms are within 4.5 Å for 75% of the trajectory. The fluctuations are defined by the expression below. Here, the reference frame 〈*R_i_*(*t*)〉 was selected as the first frame in the trajectory.

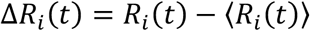

The magnitudes of these fluctuations were binned. Previous work shows that the optimal number of bins *n_opt_* is proportional to the logarithm of the number of nodes *N* in the network and can be calculated with the following expression describing the distribution of the magnitude of fluctuations as a universal feature of globular proteins^33^.

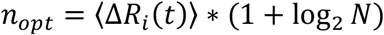

### Computing transfer entropy

From the binned timeseries, transfer entropy was computed using the expression below.

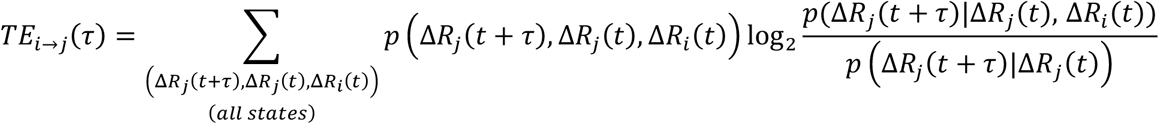

This expression describes the reduction in uncertainty of the future fluctuations of residue *j* gained by knowing the current motions of *both* residues *i* and *j* compared to only knowing those of residue *j*. Since each timeseries has its own associated noise, this noise is calculated by shuffling the binned values of timeseries *i* and re-computing the transfer entropy for a large number of trials (here, `N_shuff` = 100). This value is then subtracted from the original transfer entropy to produce an effective transfer entropy. We demonstrate the sensitivity of our computed transfer entropy to the number of shuffled trials `N_shuff` **(Fig. S1)** and its reproducibility by generating three replicas of the TE-weighted network that differ only in the shuffled TE values sampled (Supplemental Datasets).

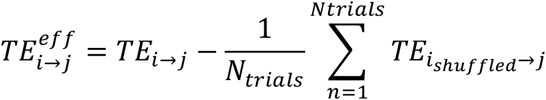

The effective transfer entropy is then normalized by the entropy rate ℎ_j_, the maximum amount information contained in timeseries *j*.

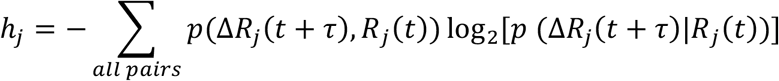

This produces a normalized transfer entropy given by the expression below.

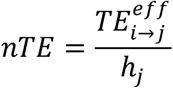

As the expression is written in the literature, *τ* would be computed by sliding a tau-length window along a trajectory written out at an output frequency assumed to be less than the value of *τ*. Using an MD simulation output frequency of 2 ps, the calculation becomes unfeasible for long trajectories. However, we are interested in these small timescales, since information between pairs of residues is maximized around 5 ps^25^. To address this discrepancy, we introduce the variable *τ*\*, which is equal to the output frequency with *τ* = 1. This allows us to assess time-lagged correlations at a wider range of timescales. The value of *τ*\* could be selected in a principled way by computing the mean and per-residue autocorrelation times from the MD trajectory and selecting a *τ*\* value that captures the timescale at which the dynamics of interest occur. To demonstrate the range of dynamic timescales within a single protein, we plotted the distribution of per-residue autocorrelation times (**Fig. S2**). For this manuscript, we primarily focus on τ* values in the high-density part of these distributions to capture directional information flow between the greatest number of residues. While we select *τ*\* = 100 ps values for generating figures in the main text, values of *τ*\* ranging from 2 ps to 5 ns have been used to capture information transfer in proteins^15,25^ and are featured throughout the supplemental material for this manuscript (Supplemental Information and Supplemental Datasets).

To assess the net direction of information flow between residues *i* and *j*, the difference between forward and reverse normalized transfer entropy values was computed as the net transfer entropy *D_i_*_→j_.

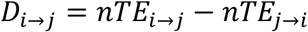

The results of these calculations are stored as NetworkX graph objects. The function `construct_causal_network` outputs two graph objects: (1) `G_causal`, a bidirectional graph which contains a forward and reverse edge for each pair of interacting residues and (2) `G_causal_net`, a directional graph containing one edge between each pair of residues with the value and direction of the net transfer entropy between them. For each edge in the `G_causal` graph object, additional values of `TE_eff`, `TE`, `maximal_TE`, and `TE_shuff` can be retrieved if the `all_vals=True` (default) parameter is passed to the `construct_causal_network` function. Due to floating-point rounding errors associated with the probability calculation, a tolerance of 0.00001 is introduced below which `TE_eff`, `TE_maximal`, and `normalized_TE` are set to zero. To improve computational efficiency, these calculations were accelerated using Numba^34^.

### Equilibrium molecular dynamics simulations

All-atom equilibrium molecular dynamics simulations were performed for checkpoint kinase 1 (Chk1, PDB ID: 1IA8^35^), TEM-1 *β*-lactamase (TEM, PDB ID: 1ZG4^36^), and cyclin dependent kinase 2 (CDK2, PDB ID: 3PXR^37^). MODELLER was used to add missing residues^38^. Simulation systems were constructed using CHARMM-GUI^39^ with 150 mM NaCl. Proteins were embedded in solvent boxes with the following dimensions: (103 Å × 103 Å × 103 Å) for Chk1, (92 Å × 92 Å × 92 Å) for TEM, and (98 Å × 98 Å × 98 Å) for CDK2. All simulations were equilibrated at 300 K with a timestep of 2 fs using the TIP3P water model^40^ and the CHARMM36 force field^41^ in an NVT ensemble with a series of gradually relaxing backbone and sidechain restraints. This was followed by 315,000 steps of additional NPT equilibration for which a weak 0.5 kcal mol^-1^ Å^-2^ positional restraint to minimize translational protein motion was applied to *C_α_* atoms of three residues. Then, 5 ns of unrestrained pre-production simulation was performed before data acquisition. All equilibration and pre-production simulations were performed using the NAMD simulation package^42^, and production simulations were performed with NAMD. Systems were simulated for a minimum of several hundred nanoseconds (500 ns for Chk1 and CDK2, 1000 ns for TEM) to ensure convergence of the transfer entropy calculation using the timescales of interest^15,16^. We demonstrate the convergence of the mean network nTE **(Fig. S3)** and per-edge nTE **(Fig. S4)** as a function of trajectory length. We performed an additional independent replicate each for Chk1 (400 ns) and CDK2 (300 ns). For TEM, we combine our simulation trajectory with eight additional published simulations of apo TEM to assess TE network variability across a substantial number of independent replicates (see Supplemental Methods)^43,44^.

To prepare these trajectories for analysis with *TEntroPy*, all proteins were stripped of water and ions and aligned to their backbone atoms. Alignment is especially important to ensure that the magnitude of residue fluctuations is not dominated by translational motion.

### Transfer entropy flux

The flux through each residue was computed by summing the normalized transfer entropy transmitted to and received from its neighboring residues in the protein contact network. This is equivalent to summing the normalized net transfer entropy between residue *i* and all its neighbors.

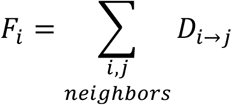

In our `NodesFlux` class, a user-defined parameter, `frac_max`, the fraction of the maximum nTE value to include as residues considered as strong broadcasters and receivers. This class also outputs a PyMOL-readable file to visualize the residues sorted as strong broadcasters (blue), weak broadcasters (light blue), weak receivers (light red), and strong receivers (red). We illustrate the robustness of the flux-based residue ranking to the choice of *τ*\*. (**Fig. S5**, Supplemental Datasets).

### Optimal path analysis

Optimal paths between pairs of source and target residues were computed using the Floyd-Warshall algorithm and reconstructed using the `floyd_warshall_predecessor_and_distance` and `reconstruct_path` functions of NetworkX^45^. This enables the comparison of paths computed from the *TEntroPy* directional graph (`G_causal`) and those computed using the MI-based network computed with dynetan.

The total entropy transfer along optimal paths is computed by summing the net transfer entropy over all pairs in the path.

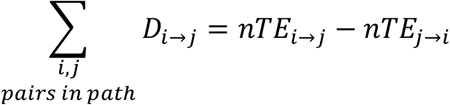

Note: Due to the overlap in the number of residues contacting both allosteric and orthosteric ligands, CDK2 was not used for optimal path and subsequent perturbation analysis.

### Studying information flow via perturbation

To mimic the decrease in residue fluctuation that occurs upon ligand binding^46^, a restraint with force constant 10 kcal mol^-1^ Å^-2^ was applied to the heavy atoms of selected residues in allosteric and orthosteric binding sites after 200 ns of production simulation. This created a 50 ns branched trajectory continued from the coordinates and velocities of the 200 ns frame, enabling direct comparison with the unrestrained trajectory. Additional replicates for Chk1 were created using a second independent MD trajectory with branch points from both the 100 ns and 200 ns frames. An additional TEM replicate was created by branching the simulated trajectory from the 300 ns frame. The similarity between replicates was quantified by comparing the per-residue fluctuation magnitudes (|Δ*R_i_*(*t*)|) and their per-edge correlations (Pearson r between |Δ*R_i_*(*t*)| and |Δ*R*_j_(*t*)|), measured relative to the mean of the 50 ns trajectory **(Fig. S6)**. We also demonstrate reasonable convergence between the per-edge correlations for the first and second halves (25 ns) of each 50 ns simulation trajectory **(Fig. S7)**. This perturbation-by-restraint approach allows us to directly access the question of how drug binding affects a protein network without performing direct binding simulations that require accurately parameterized ligands, sampling on long timescales, and extensive computational resources. Path ensembles (N=10 paths per allosteric-orthosteric residue pair) were computed directly from the `G_causal` network directly using the `shortest_simple_paths` function in NetworkX^47^.

## RESULTS

We developed and applied a workflow using our *TEntroPy* Python package to apo molecular dynamics trajectories of proteins with known allosteric and orthosteric binding sites (**Fig. 1A**). We define the residues in these binding sites as residues in contact with co-crystallized ligands. Using contact maps computed from these trajectories (see Methods), we generated directional protein networks, for which the edge weights correspond to the normalized transfer entropy. From this network, we can compute the net transfer entropy along each edge (**Fig. 1B**). We computed TE-weighted networks for checkpoint kinase 1 (Chk1), TEM-1 *β*-lactamase (TEM), and cyclin dependent kinase 2 (CDK2) to reveal directional informational flow emerging from temporal asymmetry between residue fluctuations. Structural similarity between apo and ligand-bound states of these proteins **(Fig. S8)** is consistent with the “allostery without conformational change” paradigm^48^, in which the allosteric mechanism is driven by information flow rather than large-scale conformational change.

**Fig. 1:**
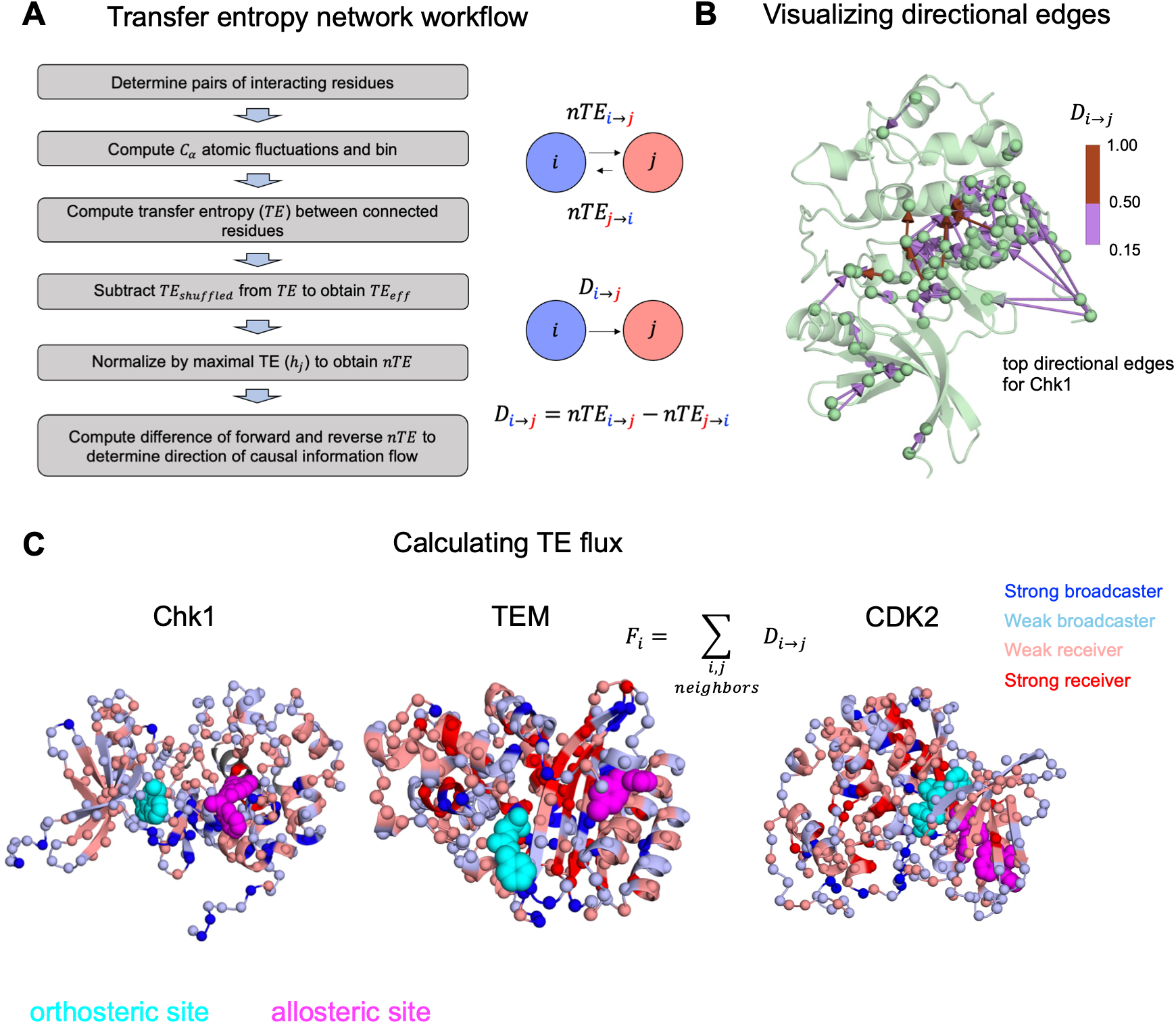
Directional protein networks were generated with the *TEntroPy* Python package. **(A)** Schematic of the workflow used for generating a directional protein network using the *TEntroPy* Python package. **(B)** Visualization of the strongest directional edges weighted by the net normalized transfer entropy (*D_i_*_→j_ = 0.15-0.50 in purple, *D_i_*_→j_ = 0.5-1 in brown) **(C)** Directional protein networks were used to compute the transfer entropy flux for each residue. Strong entropy donors generate allosteric signals and are located near important binding sites. The analysis pipeline was tested on Chk1, TEM, and CDK2.

### Transfer entropy flux reveals residues generating allosteric signals

To determine which residues in the protein network broadcast signals and which receive them, we computed the transfer entropy flux^19^ for each residue as the difference between the normalized transfer entropy transmitted and received from neighboring residues in the network. Residues are labeled broadcasters if they are high in outgoing normalized TE and receivers if they are high in incoming normalized TE. This calculation allows us to sort all residues into strong broadcasters (blue), weak broadcasters (light blue), weak receivers (light red), and strong receivers (red). We computed the transfer entropy network for a range of time lags (**Fig. S5**, see Methods).

Performing this calculation for our test proteins reveals strong broadcasters and receivers among important binding site residues. We show that these residues engage co-crystalized ligands (**Fig. 1C**). For Chk1, allosteric-site residues Ile96, Arg95, and Pro98 and orthosteric-site residues Cys87, Val68, and Leu137 were among the top entropy broadcasters, while allosteric-site residue Gly204 and orthosteric-site residue Ser147 function as the strongest receivers^49^. TEM allosteric-site residues Ile263 and Leu221 rank high among top entropy broadcasters, while Val261 ranks among top allosteric-site receivers^50^. The orthosteric site of TEM features broadcaster residues Glu240 and Ser70 and receiver residues Met69^51^. For CDK2, allosteric-site residues Leu78 and Phe80 are among the strongest receivers in the hydrophobic pocket where allosteric ligands can bind^37^. Ile35 acts as a moderate broadcaster with increased flexibility relative to hydrophobic pocket residues. Contacting ATP in the orthosteric site are strong broadcaster residue Lys129 and strong receiver residue Leu134. The presence of strong entropy broadcasters and receivers in both allosteric and orthosteric sites suggests bi-directional information flow between binding sites, in which some proteins exhibit stronger directional information flow in only one direction (allosteric site → orthosteric site), while others exhibit strong coupling in both directions (allosteric site ↔ orthosteric site). We see general agreement in residues identified as broadcasters and receivers via TE flux across independent simulation replicates (**Fig. S9**, Supplemental Datasets).

In addition to allowing us to map directional information flow onto specific structural elements of our proteins, our transfer entropy network also allows us to identify general dynamic features associated with the quantity and directionality of information. First, we compared the per-residue *C_α_* fluctuations for residues with the lowest and highest incoming and outgoing transfer entropy (**Fig. 2A-C**). Residues with high incoming TE exhibited lower average fluctuations with significantly decreased variance, indicating that these residues are more dynamically homogenous and rigid. Conversely, residues with high outgoing TE exhibited more variable, flexible dynamics. Next, we computed correlations in *C_α_* fluctuations between each pair of connected residues and compared to the total TE value (forward TE + reverse TE) for each edge (**Fig. 2D-F**). Edges with higher total TE predicted stronger pairwise correlation between residues, which demonstrates that TE captures mechanical coupling between residues.

**Fig. 2:**
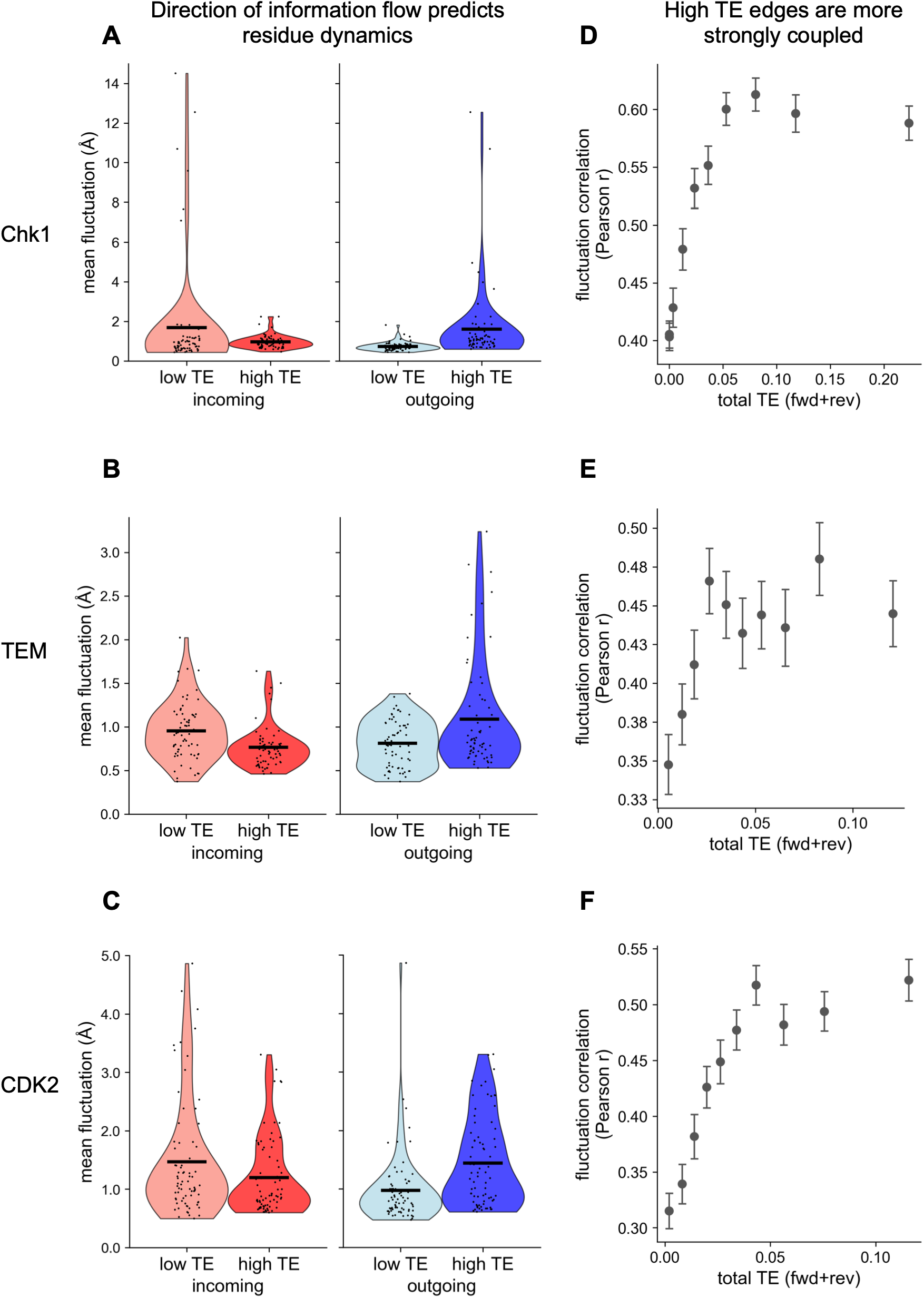
Directional information flow captures residue dynamics and pairwise coupling. **(A-C)** Comparison between the residues with the highest (top 25%) and lowest (bottom 25%) incoming (red) and outgoing (blue) mean normalized TE (nTE). Mean fluctuation magnitude distributions are shown for **(A)** Chk1 incoming (low TE residues: mean = 1.68 Å, variance = 7.85 Å^2^; high TE residues: mean = 0.97 Å, variance = 0.11 Å^2^, N = 69 residues per group) and outgoing (low TE residues: mean = 0.73 Å, variance = 0.05 Å^2^; high TE residues: mean = 1.60 Å, variance = 3.75 Å^2^, N = 69 residues per group) TE. **(B)** TEM incoming (low TE residues: mean = 0.95 Å, variance = 0.11 Å^2^; high TE residues: mean = 0.76 Å, variance = 0.06 Å^2^, N = 66 residues per group) and outgoing (low TE residues: mean = 0.81 Å, variance = 0.07 Å^2^; high TE residues: mean = 1.09 Å, variance = 0.38 Å^2^, N = 66 residues per group) TE. **(C)** CDK2 incoming (low TE residues: mean = 1.47 Å, variance = 0.97 Å^2^; high TE residues: mean = 1.20 Å, variance = 0.42 Å^2^, N = 75 residues per group) and outgoing (low TE residues: mean = 0.97 Å, variance = 0.38 Å^2^; high TE residues: mean = 1.44 Å, variance = 0.57 Å^2^, N = 75 residues per group) TE. **(D-F)** Dependence of coupling strength on the total information flow between pairs of residues (forward TE + reverse TE). All edges were divided into 10 bins. Error bars represent SEM within each bin. For each bin, the mean pairwise correlation (Pearson r) is plotted as a function of the TE value at the bin mean for **(D)** Chk1, **(E)** TEM, and **(F)** CDK2.

### Optimal paths reveal directional information flow between sites

Once we had identified residues for which changes in motion were transmitted to nearby residues, we identified pathways of directional information flow between binding sites using optimal path analysis. Using a “distance” derived from our normalized transfer entropy (see Methods), we computed paths between key broadcaster residues in one site and all residues of another binding site (**Figs. 3, S10**). We then ranked these optimal paths by maximizing difference between forward and reverse path sums. We define a parameter Δ to capture the net entropy transfer along each path. Paths with high Δ values represent dominant avenues of directional information flow between sites. A complete list of Δ values can be found in the Supplemental Datasets.

**Fig. 3:**
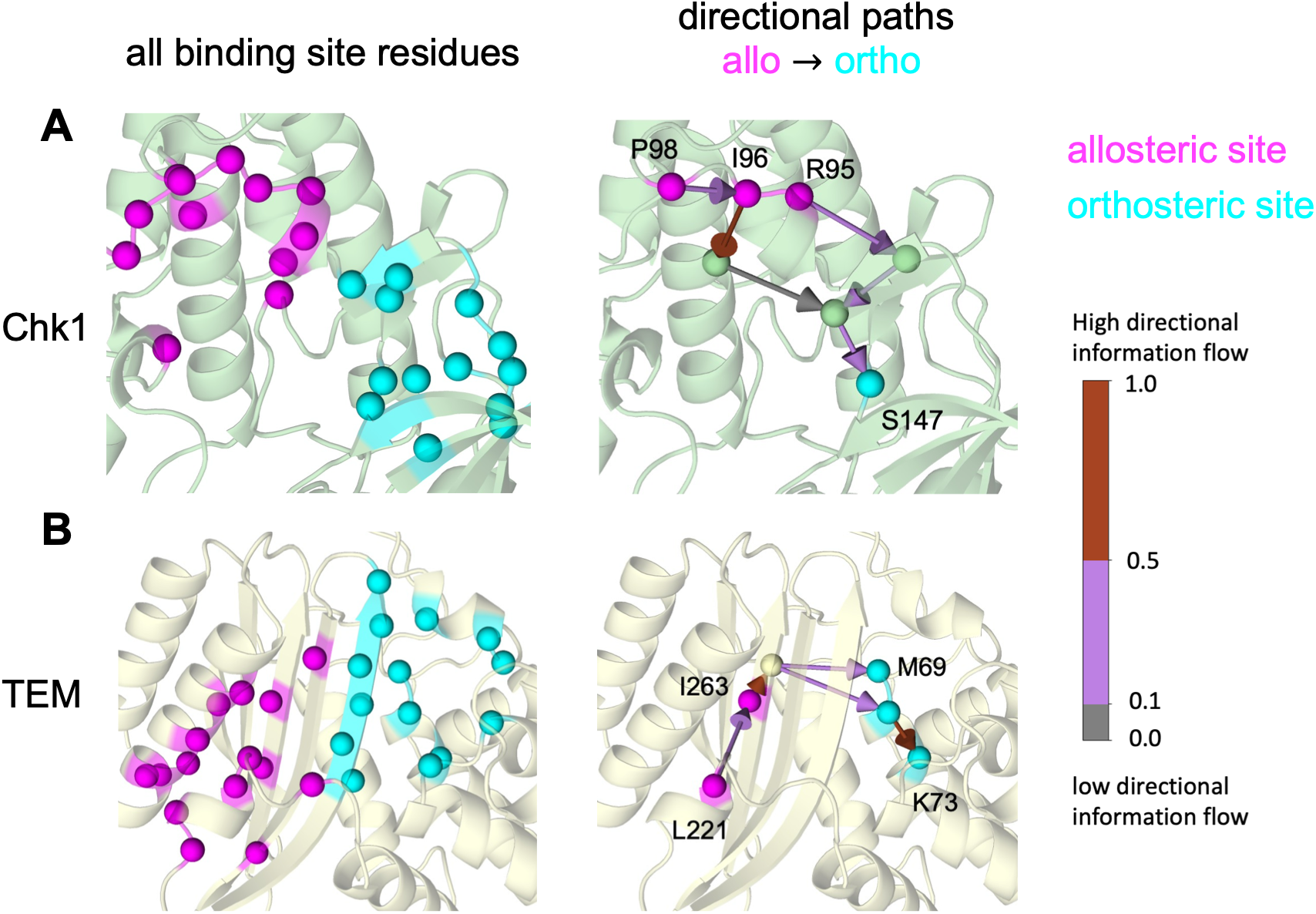
Optimal paths facilitate communication between binding sites. **(A-B)** Optimal paths were computed from allosteric to orthosteric site residues in **(A)** Chk1 and **(B)** TEM. Optimal paths were ranked by maximizing the difference between forward and reverse path sums (Δ) for each broadcaster residue in the allosteric site. Top paths are visualized using PyMOL. These high Δ value paths are the dominant avenues of directional information flow between sites. **(A)** For Chk1, top paths are shown connecting Arg95→Ser147 (Δ = 0.29), Ile96→Ser147 (Δ = 0.22), Pro98→Ser147 (Δ = 0.22). **(B)** For TEM, top paths are shown connecting Ile263→Lys73 (Δ = 0.07) and Leu221→Met69 (Δ = 0.05). A complete list of Δ values can be found in Supplemental Datasets.

For Chk1, information from allosteric site residues Ile96, Arg95, and Pro98 flows to orthosteric site residue Ser147 (**Fig. 3A**). Orthosteric sites residues Glu91, Leu84, Glu85, and Leu137 are additional reservoirs of incoming allosteric signals. In the allosteric site, residues Tyr173, Leu206, Glu205, and Pro133 receive signals from strong broadcasters in the orthosteric site **(Fig. S10A)**. Interestingly, Leu137 seems to be involved in both signal generation and receipt. Optimal paths from broadcaster residues in TEM transmit information to Lys73 and Met69 (**Fig. 3B**). Paths from broadcasters in the orthosteric site terminate at allosteric site residues Leu286, Val261, and Ile263 **(Fig. S10B)**. The set of paths from allosteric site donor residues traverse and terminate at a more diverse set of residues.

### Directional information flow was assessed under perturbation

Our transfer-entropy weighted network allows us to identify asymmetric coupling between pairs of residues and the long-range communication pathways connecting important functional sites. However, whether this network computed from equilibrium MD trajectories can predict information flow when the network is perturbed, as occurs when a drug binds to an important functional site, remains an important open question.

To study information flow between allosteric and orthosteric sites under perturbation, we applied a simple harmonic restraint of 10 kcal mol^-1^Å^-2^ to mimic the decrease in residue motion that occurs upon drug binding^46^ and measured the per-residue fluctuations for both perturbed and unperturbed simulation trajectories (**Fig. 4A**).

**Fig. 4:**
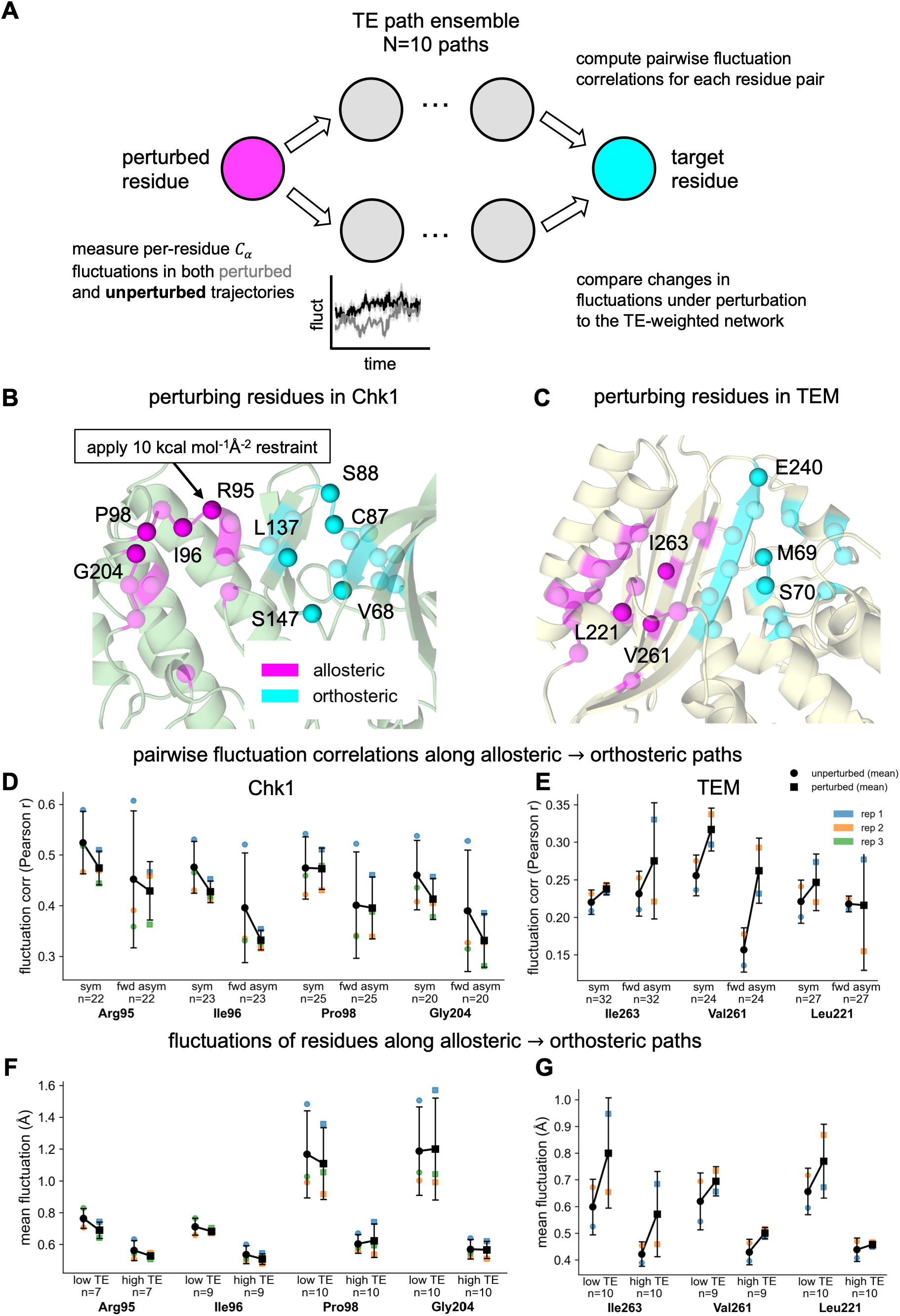
Studying directional information flow between binding sites with direct residue perturbations. **(A)** Analysis of per-residue fluctuation magnitudes and pairwise fluctuation correlations along paths from the allosteric to orthosteric site in the presence and absence of perturbation. For each perturbed residue, an ensemble of 10 paths was generated between that residue and all residues in the opposite binding site. Residue pairs in this path ensemble with the highest forward TE asymmetry (TE_forward_ – TE_reverse_, top 25%) and near-zero TE asymmetry (|TE_forward_ – TE_reverse_| ≈ 0, bottom 25%) were analyzed in both the presence and absence of perturbation. **(B-C)** Individual residues were perturbed using a 10 kcal mol^-1^Å^-2^ heavy-atom restraint. Our simulation dataset includes nine perturbed residues for **(B)** Chk1 (four allosteric site residues and five orthosteric site residues) and six perturbed residues for **(C)** TEM (three allosteric site residues and three orthosteric site residues). **(D-E)** Pairwise fluctuation correlations were computed for edges with high forward asymmetry (fwd asym) and near-zero asymmetry (sym) for each perturbation (perturbed residue name shown in bold). Error bars represent the sample standard deviation across replicates for residue pairs within each group. Differences between the mean across replicates for the unperturbed (black circles) and perturbed (black squares) dynamics were compared. Mean correlations within each bin are shown for rep 1 (blue), rep 2 (orange), and rep 3 (green). **(D)** Pairwise fluctuation correlations are plotted for forward asymmetric and near-symmetric groups of residue pairs for Chk1. **(E)** Pairwise fluctuation correlations are plotted for forward asymmetric and near-symmetric groups of residue pairs for TEM. (**F-G)** Mean fluctuation magnitudes are plotted for both high TE (top 25% of path residues) and low TE (bottom 25% of path residues) from unperturbed (black circles) and perturbed (black squares) trajectories. Error bars represent the sample standard deviation across replicates for residues in the low and high TE groups. Mean fluctuations within each bin are shown for rep 1 (blue) and rep 2 (orange). Fluctuation analysis is shown for residues on paths from allosteric perturbations to orthosteric site residues for **(F)** Chk1 and **(G)** TEM.

We performed single-site perturbation simulations (50 ns each) of nine perturbed residues in Chk1 (four allosteric site residues and five orthosteric site residues) and six residues in TEM (three allosteric site residues and three orthosteric site residues) (**Fig. 4B,C**).

We used our TE-weighted network to generate an ensemble of shortest directional paths from each perturbed residue to all residues in the opposite binding site (allosteric → orthosteric or orthosteric → allosteric). For each residue in the path ensemble, we computed the per-residue fluctuations in both perturbed and unperturbed trajectories. We also computed pairwise correlations between residue fluctuations (Pearson r) for each edge along the directional paths. We then sorted these residue pairs by their net forward TE along the path (TE_forward_ – TE_reverse_) to identify residues with high directional information flow toward the opposite binding site. We also sorted residue pairs by the magnitude of this difference to identify residues with low directional information flow (|TE_forward_ – TE_reverse_| ≈ 0) between sites.

Specifically, we compared fluctuation correlations between residue pairs with the highest forward asymmetry and near-zero forward asymmetry (TE-symmetric) for both perturbed and unperturbed trajectories (**Fig. 4D,E**). Our results reveal that both TE- symmetric and TE-asymmetric residue pairs reflect changes upon perturbation for the timescales captured by our simulations and the time lags used to compute TE-weighted networks.

How, then, might directional information flow present in the context of inter-site communication? For both Chk1 and TEM, residues along optimal paths with high incoming TE show increased rigidity (**Fig. 4F,G**). Information from neighboring residues is more easily detectable in residues with low mean fluctuations, so these residues are able to reflect a more reliable TE signal. This supports the general trend identified at the full network level (**Fig. 2**) and suggests that directional information flow can be used to identify residues involved in the interplay between rigidity and flexibility that underlies allosteric communication in proteins.

The ways in which these path residues respond to and transmit the perturbation differ across proteins and provide important insight into the proteins’ underlying allosteric mechanisms. For Chk1, the perturbing restraint generally decreased mean fluctuations in residues along optimal paths (**Fig. 4D,F and Fig. S11A,C**). Perturbing allosteric residues in Chk1 decreases pairwise correlations along optimal paths to the orthosteric site (**Fig. 4D**). Our perturbation simulations for Chk1 suggest that signals are transmitted from allosteric to orthosteric sites via mechanical rigidity, resembling allosteric mechanisms reported for other kinases in the cyclin-dependent kinase (CDK) family^52^ and in the second PDZ domain of the human protein tyrosine phosphatase 1E (hPTP1E)^53^.

In contrast, perturbing key residues in TEM enhanced both mean per-residue fluctuations and the mean pairwise correlation along optimal paths (**Fig. 4E,G and Fig. S11B,D)**. Enhanced fluctuations in the perturbed state are consistent with previous studies of TEM, in which both equilibrium simulations of the inhibitor-bound state and nonequilibrium perturbation simulations of the apo state show stronger pairwise correlations and greater per-residue fluctuations^54^.

Our perturbation results highlight both general and protein-specific mechanistic features of allosteric communication. Computing a TE-weighted network allows us to separate important residue pairs by the temporal asymmetry of their coupling, informing which residues might respond to perturbation in a time-resolved manner at a given timescale. Studying the asymmetry that emerges at small timescales could be leveraged for the development of therapeutic interventions.

## DISCUSSION

In this study, we used our *TEntroPy* Python package to build directional protein networks from molecular dynamics trajectories and used them to follow information flow from important signal-generating residues of one binding site to residues in another.

Identifying allosteric trigger residues facilitates the design of allosteric modulator drugs that would best transmit the signal to the orthosteric site and alter function. This can also allow us to identify mutations that have an asymmetric effect on inter-site communication. While the measured perturbation responses are modest for single-residue perturbations, simulations that perturb residues at multiple sites might produce responses comparable to those observed upon drug binding, where the drug engages multiple residues simultaneously. Insight gained by using our directional network to study perturbation response further demonstrates its value for studying the allosteric effects of drug binding.

Combining our TE-weighted network with perturbation simulations helps elucidate the link between the information theoretic measures and mechanical signal transmission. Our work connects the direction and magnitude of information flow to the per-residue dynamic variability and mechanical coupling of residue pairs, highlighting the interplay between rigidity and flexibility in signal transmission required for protein allostery. An important direction for future work is to integrate information metrics with existing tools for studying the mechanical responses of proteins under perturbation, such as perturbation response scanning (PRS)^55^ and strain analysis^56^.

Going beyond direct analysis of the TE-weighted network for a protein of interest, future studies could explore using a TE network to constrain generative dynamic models, such as recurrent network models (RNNs). Previous work using recurrent architectures, such as a long short-term memory (LSTM) networks, has demonstrated the link between path entropy and dynamics^57^. Transfer entropy specifically has also been used to impose feedback connections onto a feedforward deep neural network to improve model performance^58^. RNNs constrained by pairwise TE computed for multiple time lags could reveal coordinated dynamics across multiple timescales, similar to methods like time-lagged independent component analysis (tICA) used to identify slow conformational dynamics in proteins^59^. Dynamical systems analyses of these RNNs may even reveal important mechanistic insight into a protein’s dynamic stability.

By leveraging time-asymmetric directional information flow, dynamical network models, and perturbation analyses, we present a comprehensive workflow for studying allosteric signal transmission in proteins, with broad applications to the development and characterization of novel therapeutics.

## CODE AND DATA AVAILABITY STATEMENT

The source code for the *TEntroPy* Python package is deposited at the following GitHub repository: https://github.com/ryovanno/TEntroPy. MD simulation trajectories are deposited in Zenodo with DOI: 10.5281/zenodo.20777461.

## AUTHOR CONTRIBUTIONS

R.A.Y. and A.Y.L. designed the research; R.A.Y. developed the Python library, conducted molecular dynamics simulations, and analyzed the results; R.A.Y. and A.Y.L. wrote the manuscript.

## ACKNOWLEDGEMENTS

We used resources provided by the Maryland Advanced Research Computing Center (MARCC) and Advanced Research Computing at Hopkins (ARCH) at Johns Hopkins University. This work was supported by NIH grant T32GM135131 (to R.A.Y.).

