## Supplemental Information for "Directional information flow as a tool for analyzing protein allostery"

Supplemental information includes:

Supplemental Methods

Figures S1-S11

Additional data tables can be found in Supplemental Datasets (TE_supplemental_data_tables.xlsx)

### **Supplemental Methods**

*Analysis of published TEM trajectories with TEntroPy*

Additional molecular dynamics simulation trajectories of TEM in the apo form were obtained from previously published work[1,2]. In their study, initial structures for simulation were modeled from the following PDB IDs: 1ERM, 1ERO, 1ERQ, 1HTZ, 1JWP, 1LHY, 1XPB, and 3JYI. All crystal structures were mutated to the sequence of wild-type TEM-1 and co-crystallized ligands were removed. Trajectory files consist of $C_\alpha$ coordinates written out at 50 ps intervals, setting the smallest $\tau$* value for which we can compute a TE-weighted network. The authors simulated each system for 500 ns, but they removed the first 100 ns for further equilibration, resulting in 400 ns trajectories for analysis.

To make the published $C_\alpha$ trajectories compatible with the *TEntroPy* analysis pipeline, we extract $C_\alpha$ coordinates from the published text files as an array of shape (n_frames, n_res, 3), from which we generate the binned fluctuation timeseries. A single contact map generated from our 1000 ns simulation was used to ensure comparison across the same edge for all replicates.

We computed the network mean nTE and SEM across all nine trajectories (using the first 400 ns of our TEM simulation) for multiple values of $\tau$* ($\tau$* = 50 ps: 0.0103 ± 0.00158, $\tau$* = 100 ps: 0.0099 ± 0.00160, $\tau$* = 1 ns: 0.0092 ± 0.00129).

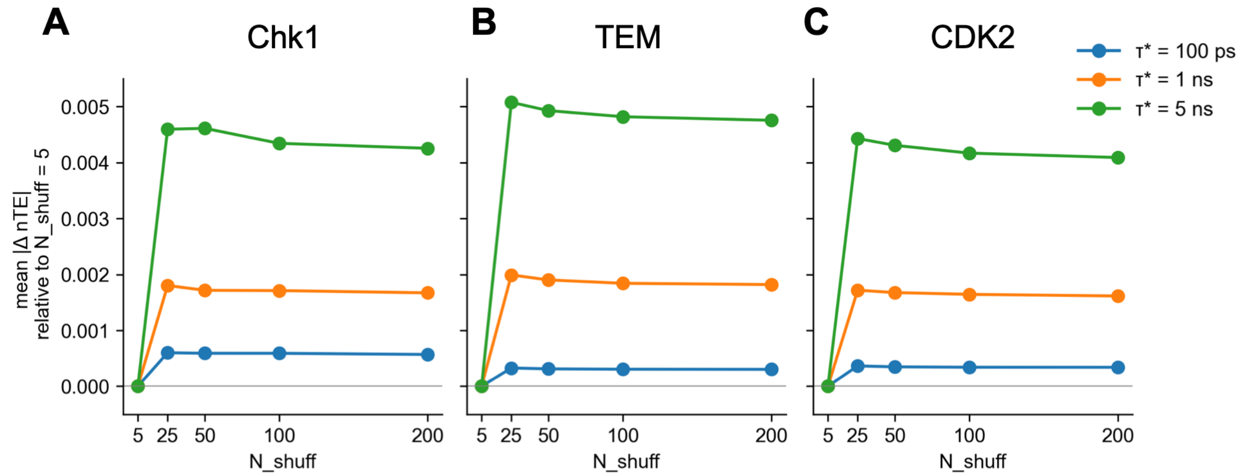


**Fig. S1: Effect of the number of shuffled trials on the mean normalized transfer entropy (nTE).** The mean absolute per-edge change in nTE (mean |ΔnTE| relative to a network computed using N_shuff = 5) was computed for different values of N_shuff = 5, 25, 50, 100, and 200. Analysis was performed on 500 ns of simulation time for $\tau^*$ = 100 ps, 1 ns, and 5 ns of proteins **(A)** Chk1, **(B)** TEM, and **(C)** CDK2.

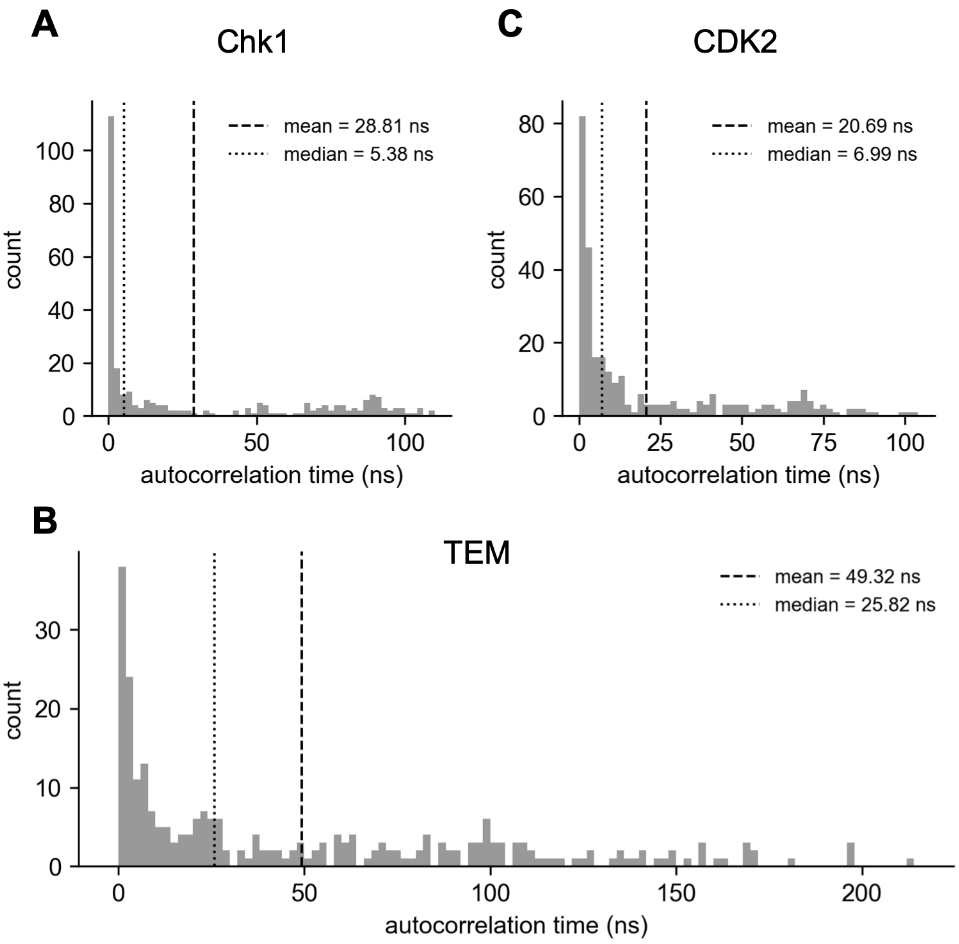


**Fig. S2: Distribution of per-residue autocorrelation times.** Per-residue autocorrelation times were computed from $C_\alpha$ fluctuations for each protein, reporting the time it takes for the autocorrelation to decay to 1/e. We plot the distribution of per-residue autocorrelation times for **(A)** Chk1, **(B)** TEM, and **(C)** CDK2 computed from the entire available simulation trajectory (500 ns for Chk1 and CDK2 and 1000 ns for TEM). The width of each histogram bin is 2 ns.

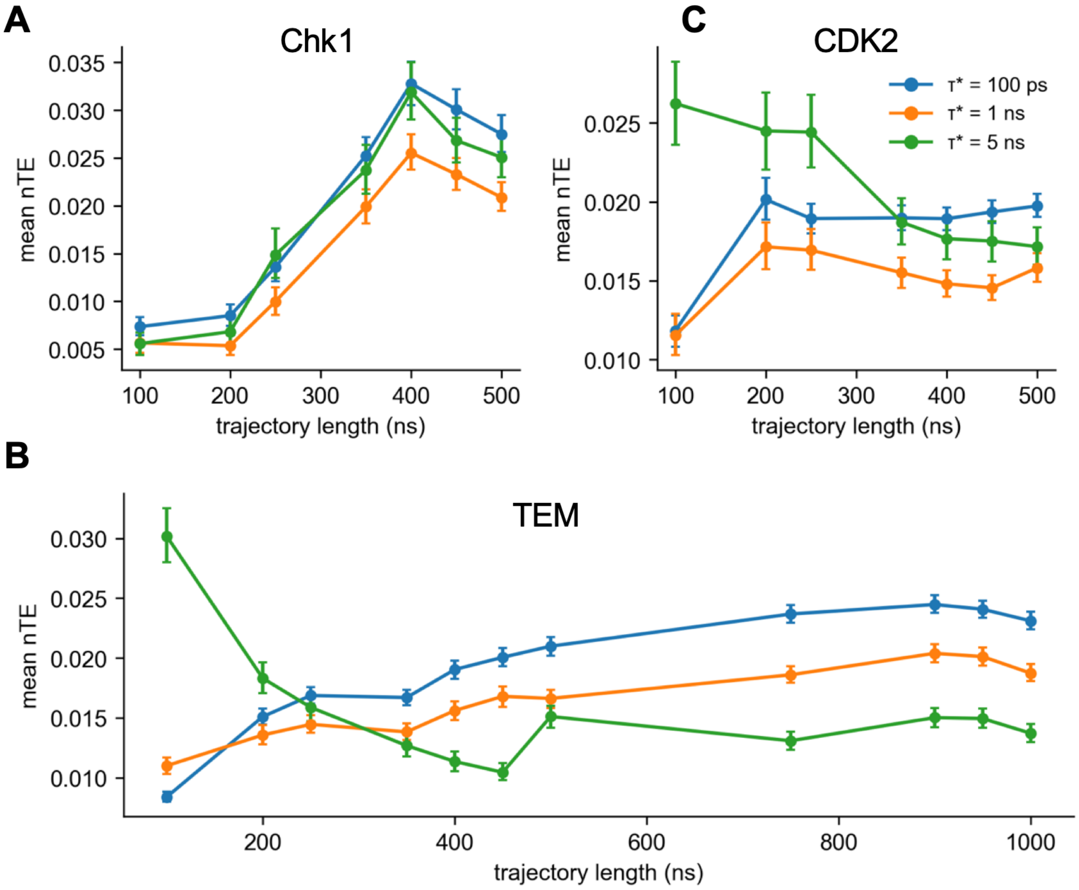


**Fig. S3: Effect of the trajectory length on the mean normalized transfer entropy (nTE).** The mean nTE was computed as a function of trajectory length using $\tau^*$ = 100 ps, 1 ns, and 5 ns for **(A)** Chk1, **(B)** TEM, and **(C)** CDK2. Error bars represent the 95% CI across edges.

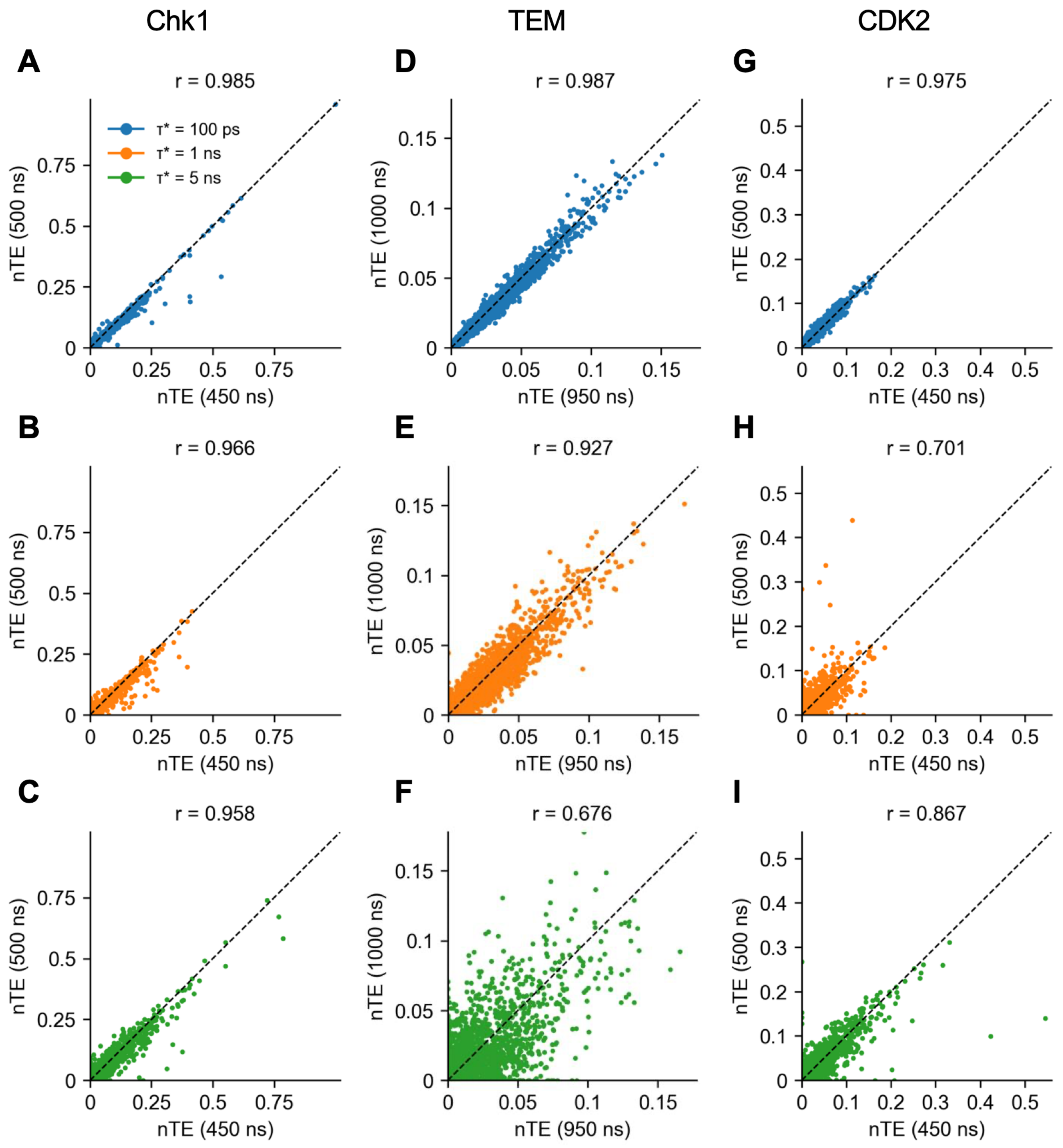


**Fig. S4: Agreement in the per-edge nTE values between the last 50 ns of simulation.** The per-edge nTE was computed from each of the last two 50 ns segments of the trajectory to illustrate convergence on the per-edge level. Pearson correlation (r) is reported for each value of $\tau^*$ for each protein simulated: **(A-C)** Chk1 (450 ns vs. 500 ns) **(A)** $\tau^*$ = 100 ps, **(B)** $\tau^*$ = 1 ns, **(C)** $\tau^*$ = 5 ns, **(D-F)** TEM (950 ns vs. 1000 ns) **(D)** $\tau^*$ = 100 ps, **(E)** $\tau^*$ = 1 ns, **(F)** $\tau^*$ = 5 ns, **(G-I)** CDK2 (450 ns vs. 500 ns) **(G)** $\tau^*$ = 100 ps, **(H)** $\tau^*$ = 1 ns, **(I)** $\tau^*$ = 5 ns.

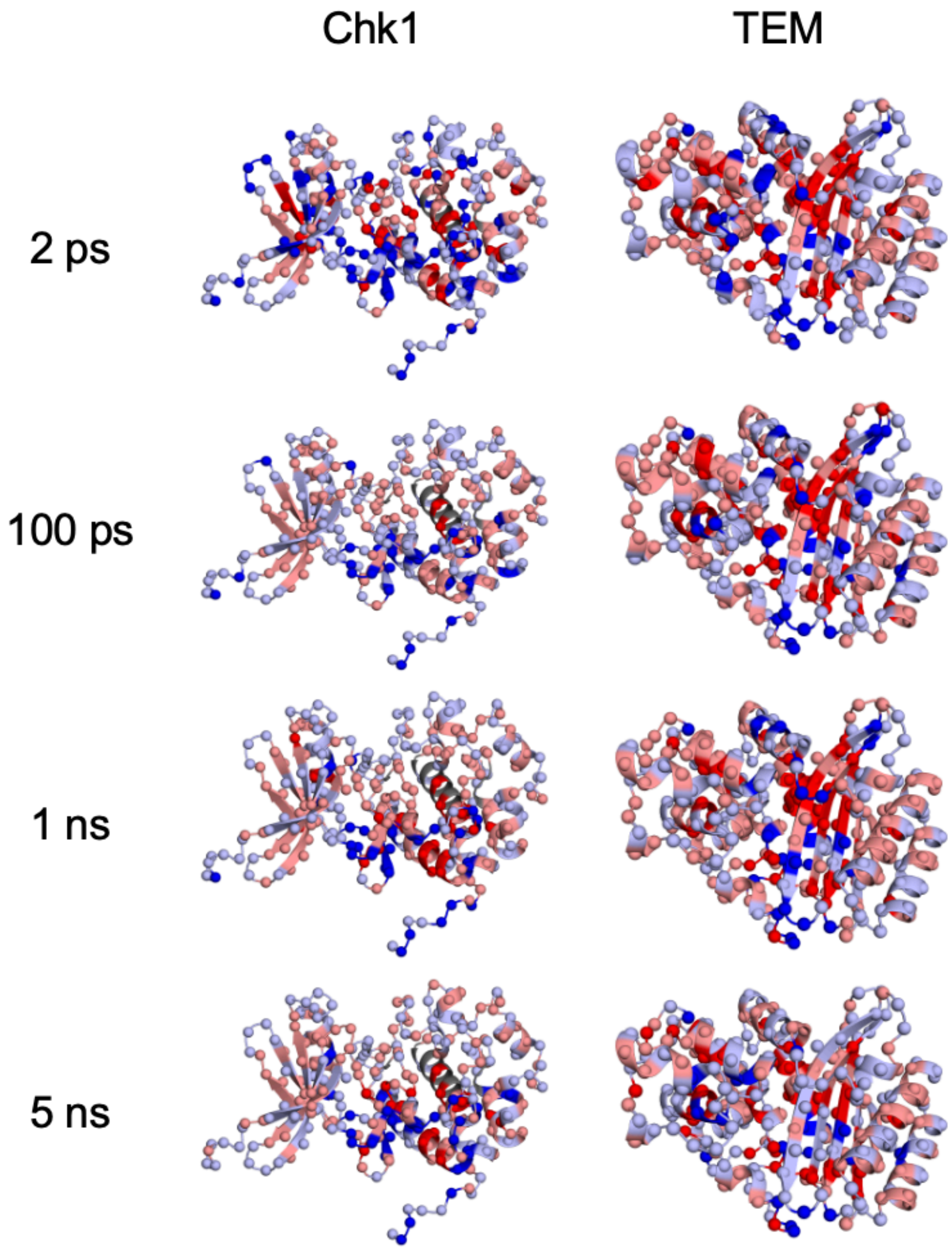


**Fig. S5:** Effect of the selection of time lag $\tau$* on transfer entropy flux. Transfer entropy flux (TE flux) was visualized for a range of different time lags: 2 ps, 100 ps, 1 ns, and 5 ns.

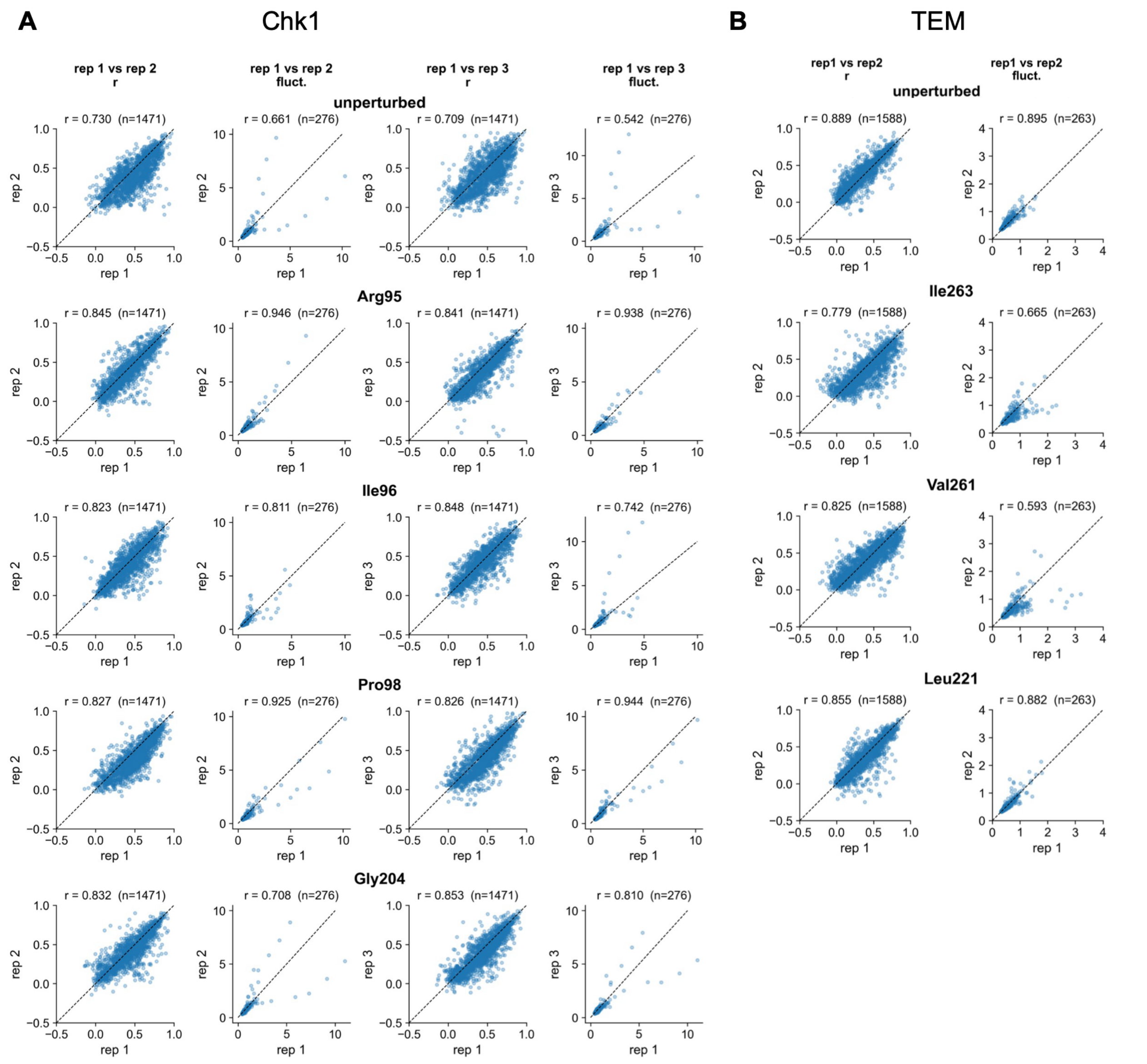


**Fig. S6: Agreement between replicate perturbation simulations for Chk1 and TEM. (A)** For Chk1, the first replicate, rep 1, represents a 50 ns simulation from a branch point at 200 ns. For Chk1, additional replicates, rep 2 and rep 3, were created from a second independent trajectory branched at 200 ns and 100 ns, respectively. The magnitude of per-residue fluctuations and pairwise correlations in fluctuation magnitude between pairs of connected residues were computed for each replicate, and Pearson correlation was computed between pairs of replicates. Comparisons are shown between replicates for the unperturbed trajectory and allosteric perturbations Arg95, Ile96, Pro98, and Gly204. **(B)** For TEM, rep 1 represents a 50 ns simulation from a branch point at 200 ns. An additional replicate, rep 2, was created from the same trajectory branched at 300 ns. Comparisons are shown between replicates for the unperturbed trajectory and allosteric perturbations Ile263, Val261, and Leu221.

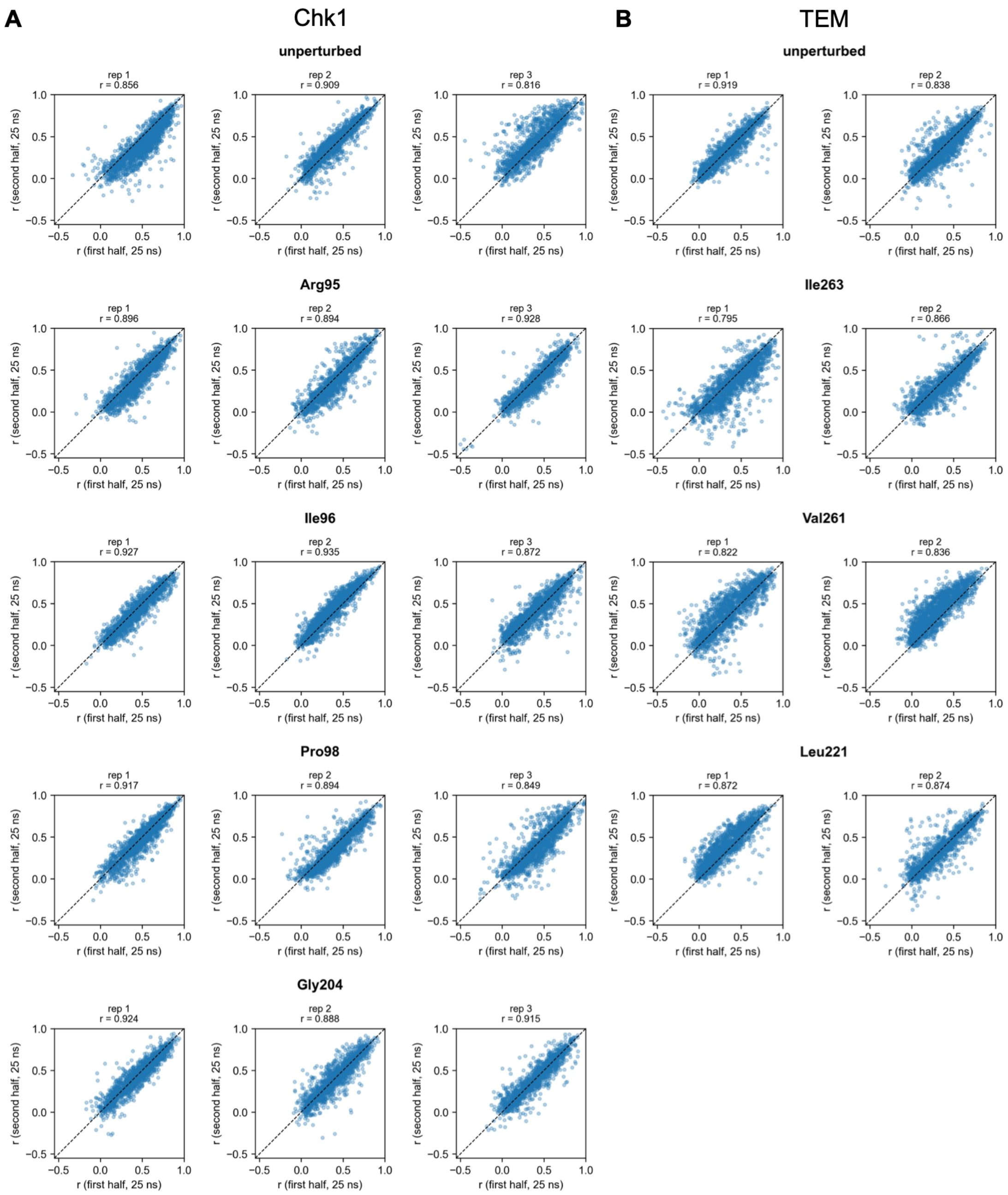


**Fig. S7: Agreement between 25 ns halves of the 50 ns perturbation simulations for Chk1 and TEM.** Correlations in per-residue fluctuation magnitude between pairs of connected residues were computed for the first and second halves (25 ns) of each simulation replicate, and Pearson r was used to quantify agreement between the two halves. **(A)** For Chk1, intra-replicate agreement is plotted for the unperturbed trajectory and allosteric perturbations Arg95, Ile96, Pro98, and Gly204 **(B)** For TEM, intra-replicate agreement is plotted for the unperturbed trajectory and allosteric perturbations Ile263, Val261, and Leu221.

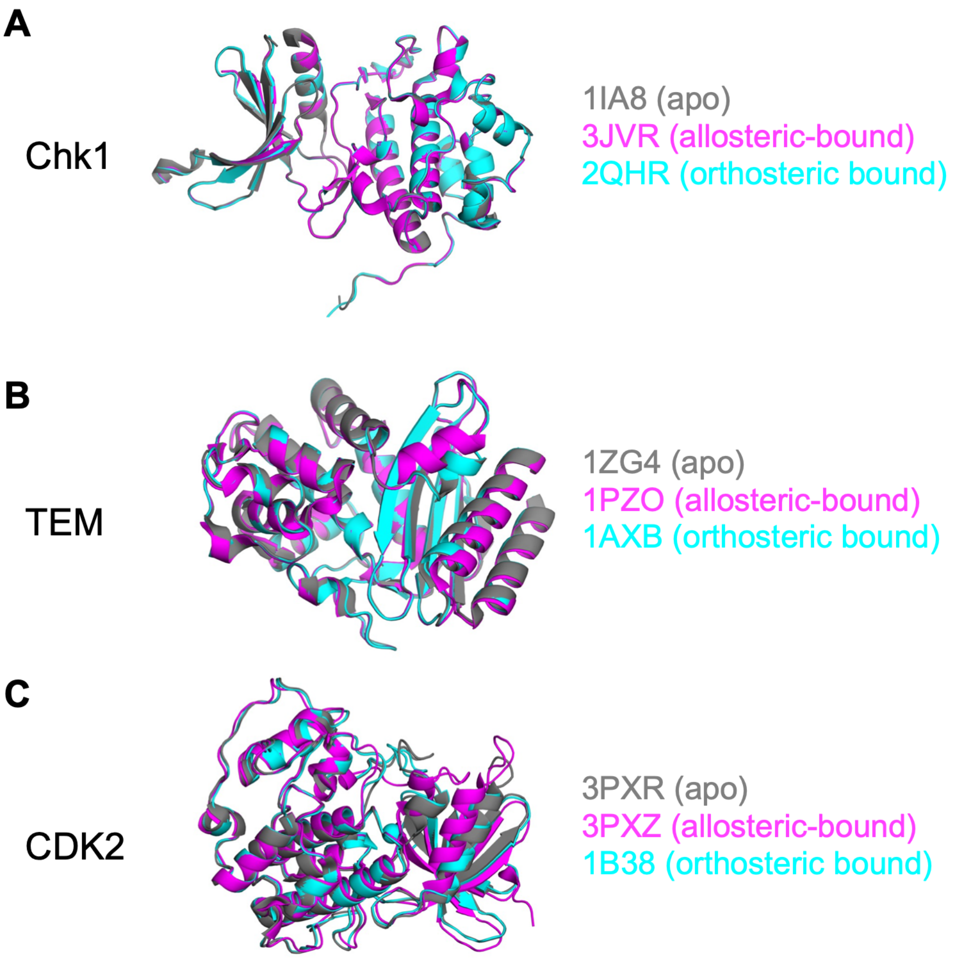


**Fig. S8: Similarity between apo, allosteric-bound, and orthosteric-bound crystal structures of Chk1, TEM, and CDK2.** Apo (gray), allosteric-bound (magenta), and orthosteric-bound (cyan) crystal structures are shown for each protein discussed in this manuscript. Holo structures are backbone-aligned to the apo structure, and backbone RMSD is reported relative to the apo structure. PDB IDs for each structure are included. **(A)** Chk1: 1IA8 (apo), 3JVR (allosteric-bound, RMSD = 0.343 Å), 2QHN (orthosteric-bound, RMSD = 0.499 Å), **(B)** TEM: 1ZG4 (apo), 1PZO (allosteric-bound, RMSD = 0.881 Å), and 1AXB (orthosteric-bound, RMSD = 0.273 Å), **(C)** CDK2: 3PXR (apo), 3PXZ (allosteric-bound, RMSD = 2.307 Å), and 1B38 (orthosteric-bound, RMSD = 0.687 Å).

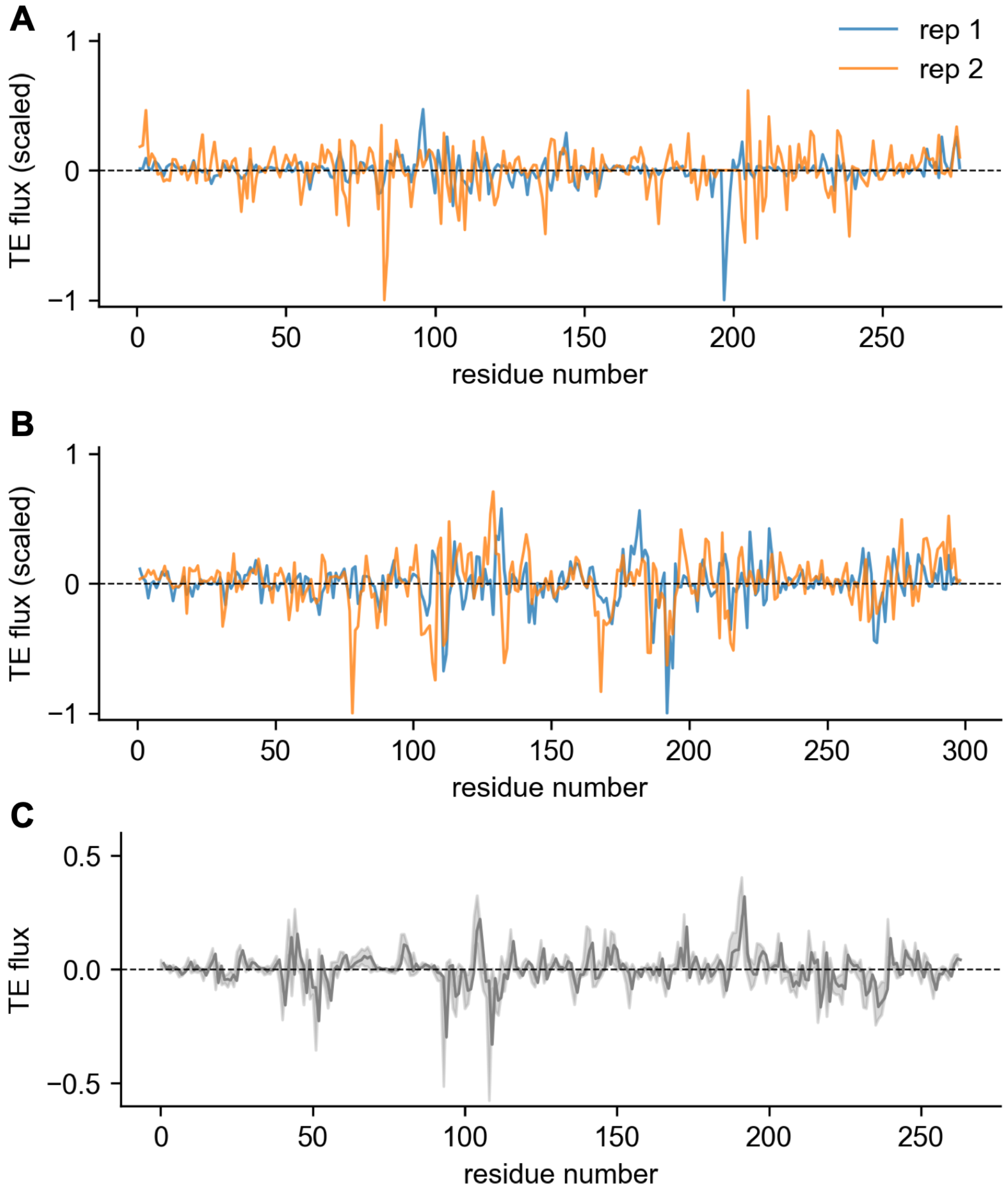


**Fig. S9: Comparison of TE flux across independent simulation replicates. (A)** TE flux (scaled by the maximum TE flux value for each replicate) is shown for two independent simulations of Chk1. The TE networks for both replicates were calculated from 400 ns MD trajectories using $\tau^*$ = 100 ps (Spearman $\rho$ = 0.16). The purpose of scaling is to demonstrate our ability to identify residues acting as broadcasters and receivers across replicates via ranked TE flux. **(B)** Scaled TE flux is shown for two independent simulations of CDK2. The TE networks for both replicates were calculated from 300 ns MD trajectories using $\tau^*$ = 100 ps (Spearman $\rho$ = 0.29). **(C)** The average TE flux was computed over nine 400 ns simulations of TEM using $\tau^*$ = 100 ps. The shaded region represents SEM across replicates. Additional details comparing TE flux across replicates can be found in Supplemental Datasets.

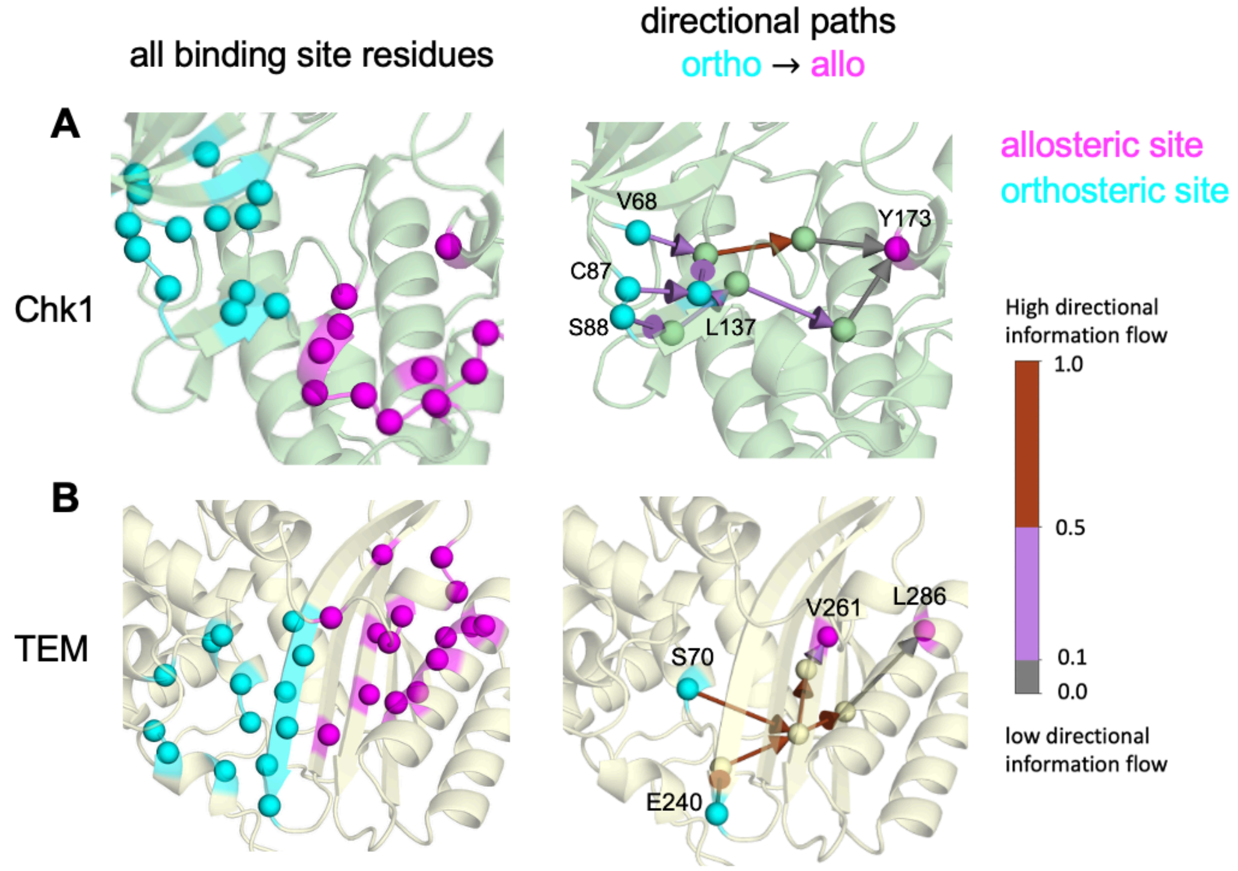


**Fig. S10:** Directional paths from orthosteric site residues to allosteric site residues in **(A)** Chk1 and **(B)** TEM. A complete list of Δ values ranking directional paths can be found in **TE_supplemental_data_tables.xlsx**.

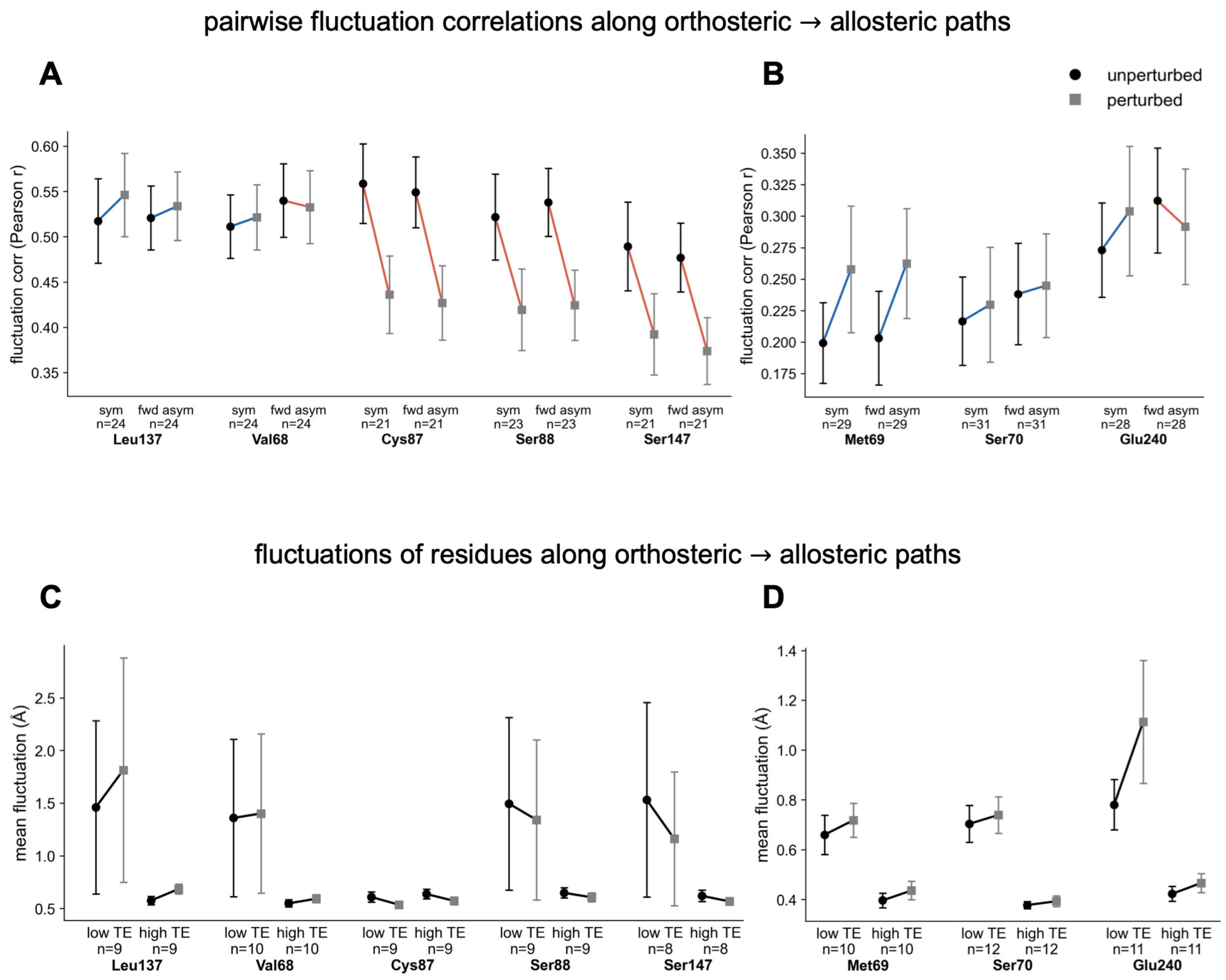


**Fig. S11: Analysis of pairwise correlations and per-residue fluctuations along paths from perturbed orthosteric residues to the allosteric site. (A-B)** Pairwise fluctuation correlations were computed for residue pairs in the path ensemble with the highest forward TE asymmetry ($TE_{forward} – TE_{reverse}$, top 25%) and near-zero TE asymmetry ($|TE_{forward} – TE_{reverse}| \approx 0$, bottom 25%) for each orthosteric site perturbation for **(A)** Chk1 and **(B)** TEM (perturbed residue name shown in bold). Error bars represent the SEM across edges within each group. Differences between unperturbed (black circles) and perturbed (gray squares) dynamics were visualized for forward asymmetric and near-symmetric pairs. Red and blue connecting lines indicate decreased or increased correlations upon perturbation, respectively. **(C-D)** Mean fluctuations are plotted for both high TE (top 25% of path residues) and low TE (bottom 25% of path residues) from unperturbed (black circles) and perturbed (gray squares) trajectories. Error bars represent the SEM across residues in the low and high TE groups. Fluctuation analysis is shown for residues on paths from orthosteric perturbations to allosteric site residues for **(C)** Chk1 and **(D)** TEM.
